# Solution State Methyl NMR Spectroscopy of Large Non-Deuterated Proteins Enabled by Deep Neural Networks

**DOI:** 10.1101/2023.09.15.557823

**Authors:** Gogulan Karunanithy, Vaibhav Kumar Shukla, D. Flemming Hansen

## Abstract

Methyl-TROSY nuclear magnetic resonance (NMR) spectroscopy is a powerful technique for characterising large biomolecules in solution. However, preparing samples for these experiments is arduous and entails deuteration, limiting its use. Here we demonstrate that NMR spectra recorded on protonated, uniformly ^13^C labelled, samples can be processed using deep neural networks to yield spectra that are of similar quality to typical deuterated methyl-TROSY spectra, potentially providing more information at a fraction of the cost. We validated the new methodology experimentally on three proteins with molecular weights in the range 42-360 kDa and further by analysing deep learning-processed NOESY spectra of Escherichia coli Malate Synthase G (81 kDa), where observed NOE cross-peaks were in good agreement with the available structure. The new method represents a substantial advance in the field of using deep learning to analyse complex magnetic resonance data and could have a major impact on the study of large biomolecules in the years to come.

## Introduction

Nuclear Magnetic Resonance (NMR) spectroscopy is a ubiquitous technique in material science, chemistry, structural biology, and in clinical diagnosis. In bioscience, NMR provides unprecedented insight into functional motions and on non-covalent interactions with atomic resolution. However, NMR is notoriously insensitive, so maximising resolution and sensitivity is a perpetual challenge within all areas of NMR spectroscopy. To this end, nuclear spin-relaxation, which is the process by which equilibrium magnetisation is restored and detectable NMR signal is lost, scales rapidly with molecular size, making it challenging to study large biomolecular systems by solution-state NMR. This has meant that individual NMR experiments are traditionally each associated with size limits of the system under investigation, above which most signals are broadened beyond detection.

Over many decades, a series of developments have raised the size-limits of detection in biomolecular applications, combining advances in hardware, sample preparation and pulse sequence development. The introduction of methyl-TROSY methods^1^, wherein methyl-bearing side chains are used to probe biomolecular structure and dynamics, provided a step-change in molecular weight limitations for solution-state biomolecular NMR. Using these techniques makes it possible to study systems up to the megadalton molecular weight range. A key requirement, however, for attaining high quality methyl-TROSY spectra is that the protein should be prepared with a very high level of deuteration. Consequently, in practice, for methyl-TROSY NMR studies, the proteins produced are completely deuterated with the exception of [^1^H, ^13^C] labelled methyl moieties in specific side chains, typically those in isoleucine, leucine, methionine, and valine. The labels are introduced using specific pre-cursor compounds that are tuned to specific side chains and robust protocols exist^2,3^. However, uniform deuteration adds several disadvantages, such as a considerable extra cost to protein production, typically lower yields of expressed protein, and it is not even possible in many systems of considerable biological interest, including proteins that can only be expressed in a mammalian system. As such, the ability to obtain high-quality ^13^C-^1^H correlations maps from uniformly ^13^C labelled protonated proteins would be highly desirable. The uniform labelling is easier, cheaper, and gives access to peaks associated with all methyl-bearing side chains rather than just those where the appropriate precursor has been added during protein expression. Such a method would also avoid the need for deuteration and pave the way for characterisations of large proteins that can only be expressed in mammalian systems.

In this current era with burgeoning applications and developments of AI, from computational structural biology^4,5^ to sophisticated large language models^6^, it is natural to look for solutions within the field of AI to the challenges encountered in characterising large proteins. In this context, it has over the last couple of years been shown by others and us that deep neural networks (DNNs) can indeed be trained to accurately transform^7–9^ and analyse^10,11^ complex NMR data. The most recent applications use supervised deep learning, where a DNN is supplied with an input and target training dataset and through a training process the DNN ‘learns’ the mapping between the two. Typically, this training requires very large amounts of training data, but importantly, as has been noted in several prior studies^8,12,13^, it is possible to simulate an arbitrary amount of realistic training data for magnetic resonance based supervised deep learning, avoiding a significant potential data bottleneck. Deep learning methods have now been successfully applied to several tasks in magnetic resonance spectroscopy including the analysis of DEER data^12^, reconstruction of non-uniformly spectra^7,9,13^, peak-picking^11,14^, and virtual homonuclear decoupling^8^.

Herein we demonstrate that deep neural networks (DNNs) can be used, in place of the traditional 1822 Fourier transform^15^, to deliver very high-quality ^13^C-^1^H correlation spectra from uniformly ^13^C protonated samples of even large proteins. In brief, the DNNs presented below are trained to map the broad ^13^C-^1^H spectra of uniformly labelled protonated samples to spectra that are akin to classical methyl-TROSY spectra. Specifically, the DNNs are applied to time domain NMR data, remove the effects of one bond ^13^C-^13^C scalar couplings in the ^13^C dimension, and increase the resolution for both the ^1^H and ^13^C dimensions by effectively sharpening the observed cross-peaks. The net effect of these two processes is that following the application of these DNNs, the appearance of peaks associated with methyl bearing side chains in protonated samples are approaching those attained from deuterated samples with specifically labelled side chains. A schematic illustration and summary of the effects of the DNNs is provided in Figure 1, where the characterisation of the 81 kDa MSG is used as an example.

**Figure 1:**
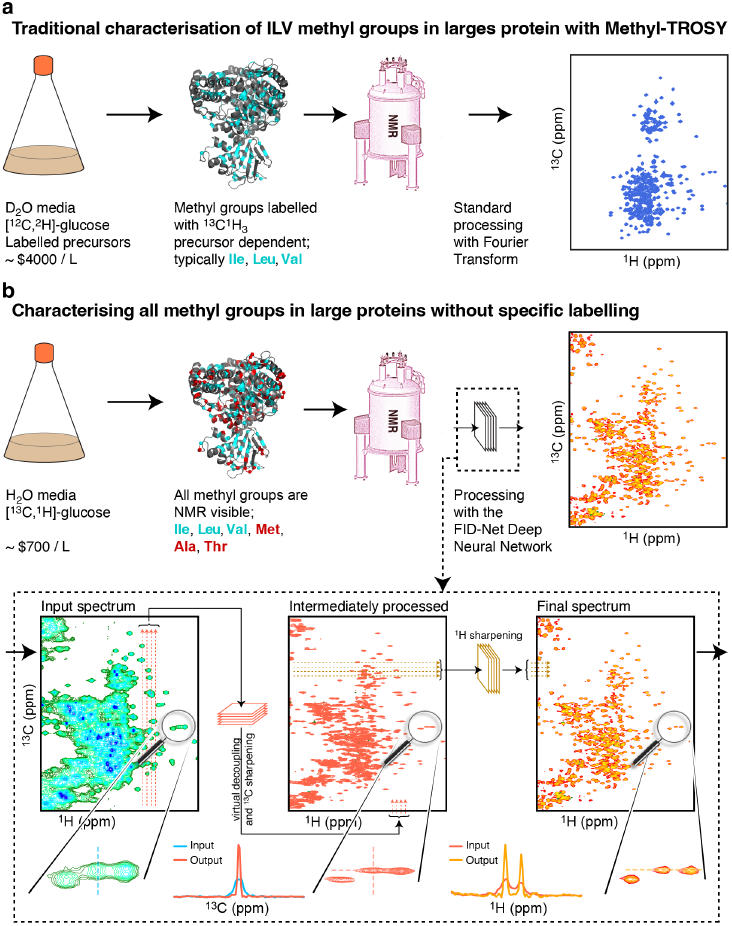
Overview of processing NMR spectra with FID-Net. **(a)** Overview of traditional tools used to characterise methyl groups in large proteins, which requires expression in bacterial cells such as *E. coli*, deuteration of the protein, and specific isotopic labelling. **(b)** Overview of our new method to characterise large, non-deuterated, uniformly labelled proteins enabled by processing with the deep neural network FID-Net. Two FID-Net networks are trained, (*i*) one network trained to virtually decouple and enhance the in the ^13^C dimension of the initial 2D ^13^C^1^H correlation spectra (green-blue to red spectra), (*ii*) followed by a second network trained to enhance the resolution in the ^1^H dimension (red to orange spectra). The example show is that of Malate Synthase G (MSG) an 80 kDa protein. As the protein is uniformly labelled it gives rise to peaks associated with all methyl groups in the protein, including methionine, alanine and threonine residues as well as isoleucine γ2 methyl groups. The additional methyl probes offered by the uniform labelling scheme are highlighted on the structure of MSG (red). The estimated costs in a and b are calculated using listed prices from Sigma-Aldrich.

We robustly cross-validate the trained DNNs on synthetic data and show the applicability of the trained DNNs on experimental data, where we demonstrate the effectiveness of this methodology on a range of increasingly large proteins: HDAC8 (42 kDa), Malate synthase G (MSG, 81 kDa) and α7α7 (360 kDa), demonstrating how the DNNs provide a highly effective route to studying large proteins by NMR. Finally, we apply the new method to obtain 3D Methyl NOESY NMR spectra of MSG, which can aid in chemical shift assignments and/or structural characterisations.

## Results

Attempting to measure high-quality ^13^C-^1^H correlations maps on large proteins using classical approaches, such as ^13^C-^1^H HSQC spectra, run into several problems. Firstly, since the proteins are uniformly ^13^C labelled they will be subject to one-bond ^13^C-^13^C scalar couplings that will evolve during indirect chemical shift evolution, splitting signals into multiplets and complicating interpretation of the spectrum. Of perhaps greater significance, is that the lack of deuteration in the system which means that signals will be significantly broadened in both the ^13^C and ^1^H dimensions as a result of significantly increased dipolar relaxation. Consequently, peaks in the spectra will be very difficult to identify and difficult to assign to specific sites in the protein making the spectra challenging to interpret and limiting the utility of such a labelling scheme. Other tools such as constant-time ^13^C-^1^H HSQC spectra^16,17^ also do not provide good spectra of large uniformly labelled proteins, since the constant-time substantially skew the intensities and even renders many of the signals invisible. However, due to the inherent sensitivity of methyl groups, ^13^C-^1^H correlations maps of protonated large proteins nonetheless contain a significant amount of information from many of the methyl groups present. The challenge is that these spectra are very hard to interpret, even by specialists, due to the poor resolution, see *e*.*g*. Figure 1b.

### Training and cross-validation on synthetic data

In order to transform ^13^C-^1^H correlation maps from universally ^13^C labelled proteins into spectra that can easily be interpreted we train two DNNs, both of which are based on the FID-Net architecture. Full training details for the DNNs are provided in the supplementary information. Briefly, the first network is trained to transform time-domain FIDs in the ^13^C dimension by removing a single cosine modulation corresponding to a ^13^C-^13^C coupling constant and reducing the decay rate of the peak such that it gives a sharper signal in frequency space. The second DNN is trained to act on FIDs in the ^1^H dimension. In this case the network is trained only to reduce the decay rate of FIDs so that peaks are sharper in the frequency domain of this dimension. Both networks are trained independently solely on synthetic data (full parameters given in the supplementary information).

For transforming a full ^13^C-^1^H 2D plane of a uniformly labelled protein the full workflow is as follows: the input 2D plane is first processed and Fourier transformed in the ^1^H dimension before being transposed. This half-processed spectrum is then passed to the first DNN where the signal modulation due to one-bond ^13^C-^13^C couplings is removed and the signals are sharpened. Subsequently the ^13^C dimension can be processed and Fourier transformed as normal. The spectra is then transposed back to the ^1^H dimension and inverse Fourier transformed and Hilbert transformed. The resulting time domain data is passed to the second DNN to sharpen signals in the ^1^H dimension. This ^1^H dimension is then reprocessed, and Fourier transformed to yield the final frequency-domain spectrum.

In order to test and benchmark this approach it is first applied to synthetic data. Rather than using randomly generated data, as is done in the training of the networks we aim to benchmark performance on synthetic data that is nonetheless reminiscent of actual ^13^C-^1^H correlation maps of proteins. In order to do this, synthetic spectra are made using chemical shifts expected for real systems as sampled from the BMRB^18^. Using this approach, we generate one hundred synthetic spectra with a comparable number of peaks to the 42 kDa HDAC8^19^ and one hundred synthetic spectra with a comparable number of peaks to MSG^20^. These spectra contain all expected ^13^C-^13^C couplings as well as broad peaks as expected for large, protonated proteins. Given that these spectra are synthetically generated we can also generate the ‘ideal’ target spectrum in which all ^13^C-^13^C scalar couplings are removed and the linewidths are narrowed and we know the positions of all peaks within this target spectrum. In order to test the performance of the DNNs we transform the original synthetic spectra using our two trained DNNs. We then pick peaks in the resulting transformed spectra and compare the results against the known peak positions looking at the rate of both true positives and false negatives. In order to avoid any influence of the accuracy of the peak-picking algorithm used, which can vary, only peaks that are isolated in the processed spectra (distance larger than processed linewidths) are considered.

As shown in Figure 2, for synthetic spectra, the FID-Net processing appears to work successfully at significantly enhancing the resolution of spectra expected for uniformly ^13^C-labelled proteins even when there are a number of large overlapping signals. Based on the high levels of true positive peaks and low levels of false positive peaks observed on synthetic data, we proceed to testing the FID-Net based processing to experimental data, where the proteins are uniformly ^13^C labelled.

**Figure 2:**
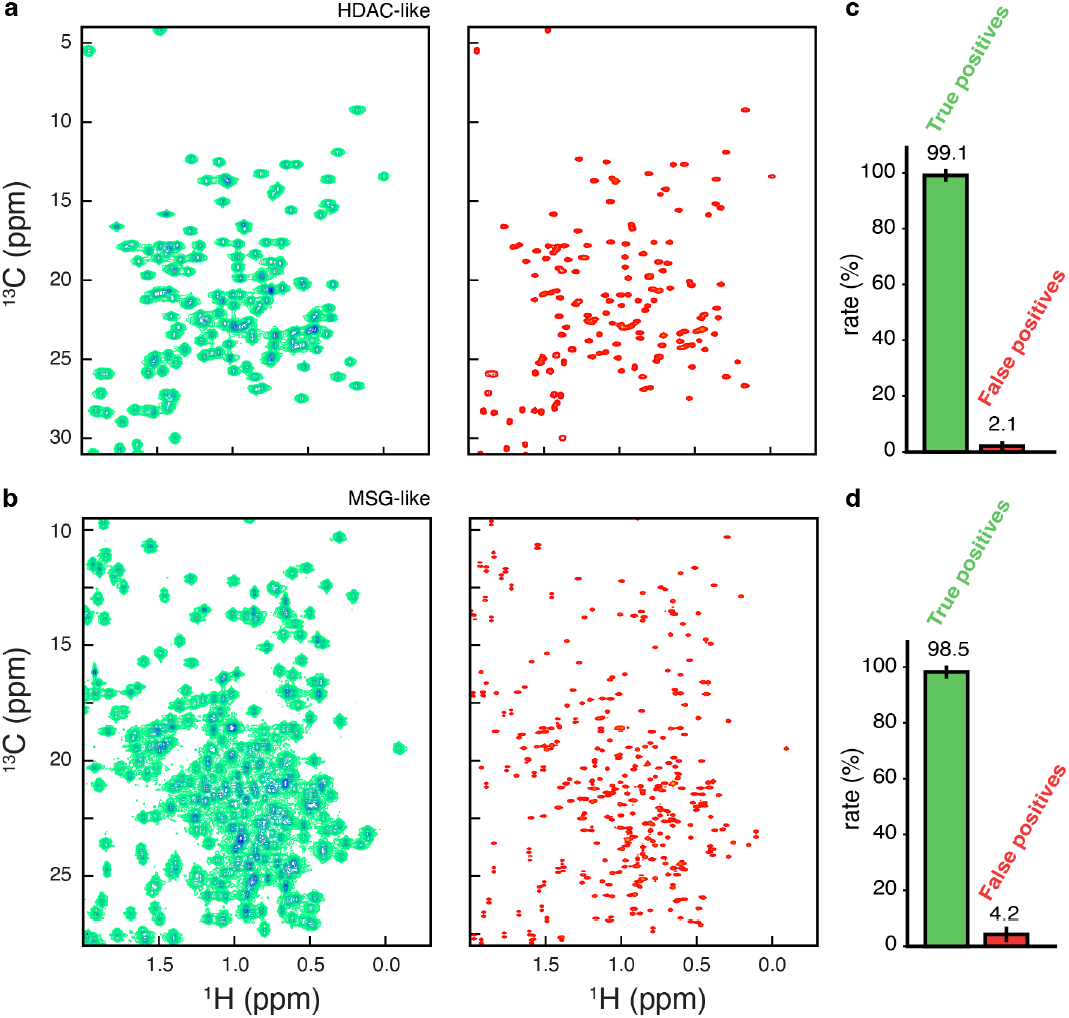
Cross-validation on synthetic data. Exemplar synthetic data **(a)** and **(b)** without analysis with the FID-Net DNNs (left column) and with FID-Net based processing right hand column. **(a)** shows a synthetic spectrum where we have a similar number of signals to HDAC8 and with **(b)** the spectrum has a similar number of signals to MSG. In both cases one hundred distinct spectra with a similar number of signals to those shown in **A** or **B** are generated. They are then analysed using the FID-Net approach and the resulting spectra are peak picked. From FID-Net analysed spectra peaks are picked and compared to ground truth values. From picked peaks true positive and false negative rates of peaks are calculated (only considering isolated peaks) and plotted in **(c)** and **(d)**. Full details are given in Supplementary Information.

### Application to experimental 2D ^13^C-^1^H correlation spectra

To test the feasibility of our approach to transform experimental ^13^C-^1^H correlation spectra of uniformly ^13^C labelled proteins the method was applied to increasingly larger proteins, demonstrating the ability of the FID-Net approach to provide high quality correlation maps, similar in quality to methyl TROSY spectra. In addition to being far less costly than deuterated analogues and typically giving higher yields during expression, a key advantage of the methodology here is that information is provided on all methyl-bearing side chains, including, alanine ^13^C^β^, isoleucine ^13^C^γ2^ and ^13^C^δ1^, leucine ^13^C^δ1,δ2^, methionine ^13^C^ε^, threonine ^13^C^γ2^, and valine ^13^C^γ1,γ2^ since the sample is uniformly ^13^C labelled. It should be mentioned that methods do exist for specific labelling of nearly all methyl groups^21^, however, these approaches typically lead to reduced yields and come with substantially larger costs.

Following cross-validations on synthetic data, the trained FID-Net networks were applied to the 42 kDa protein HDAC8^19^. While a protein of this size is relatively small for methyl-TROSY studies, as shown in the ^13^C-^1^H correlation spectrum in Figure 3a (blue-green) of a nondeuterated, uniformly ^13^C labelled sample, signals in the methyl region of the spectrum are nonetheless broad and overlapped, making it difficult to discern many of the signals. This also holds for a constant-time ^13^C-^1^H HSQC spectrum, where the constant-time (27 ms or 54 ms) substantially skew the intensities and renders many of the signals invisible, see Figure S1. Conversely, following application of the FID-Net Networks (orange spectrum), Figure 3b, the signals are virtually decoupled in the ^13^C dimension and sharpened in both the ^13^C and ^1^H dimensions. This makes peak identification within these spectra much more straightforward. By overlaying the FID-Net transformed spectrum with the classical methyl-TROSY ILVM spectrum (blue) of a deuterated sample of HDAC8, Figure 3c, where only the side chains of these amino acids are labelled, an excellent correspondence is seen between isoleucine, leucine and valine methyl peaks with the FID-Net processed spectrum. The linewidths of the peaks in both of these spectra are highly comparable and all expected peaks from the spectrum are recovered in the FID-Net processed spectrum. Additional peaks are also visible in the FID-Net processed spectrum, due to the presence of additional labelled methyl groups in the sample, such as, threonine and isoleucine ^13^C^γ2^. Small peak-shifts are mainly due to the isotope shifts originating from deuteration^22^. Full overlay of the FID-Net processed spectrum of HDAC8 and the methyl-TROSY spectrum is shown in Figure S2.

**Figure 3:**
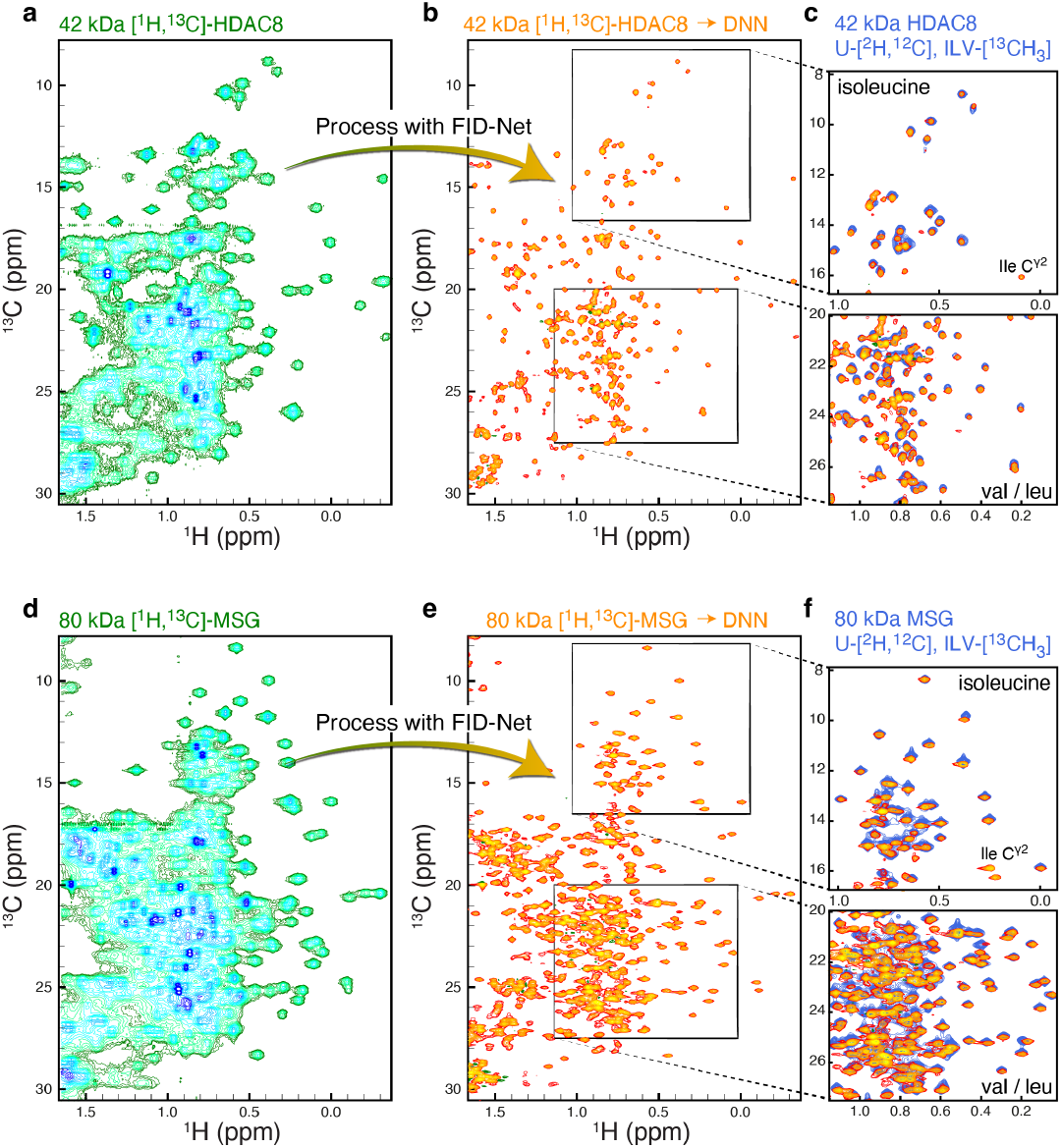
FID-Net processed methyl HSQC spectra of uniformly ^13^C labelled non-deuterated proteins. **(a)** A ^13^C-^1^H HSQC NMR spectra of uniformly ^13^C labelled, non-deuterated, HDAC8 (42 kDa) processed with a standard discrete Fourier transform. **(b)** The spectrum in a processed with the FID-Net DNNs. **(c)** Comparison of FID-Net processed HSQC spectrum in **b** (orange) with a methyl-TROSY HMQC spectrum of an ILV specifically labelled and deuterated HDAC8 (blue). **(d)** A ^13^C-^1^H HSQC NMR spectra of uniformly ^13^C labelled MSG (80 kDa) processed with a standard discrete Fourier transform. **(e)** The spectrum in d processed with the FID-Net DNNs. **(f)** Comparison of FID-Net processed HSQC spectrum in d (orange) with a methyl-TROSY HMQC spectrum of an ILV specifically labelled and deuterated MSG (blue) for two selected regions. Many methyl groups are not visible in the ILV labelled sample, such as, Isoleucine ^13^C^γ2^ (labelled).

To test the robustness of the FID-Net processing approach on larger systems, with substantially more crosspeaks and signal overlap, we next applied the FID-Net DNNs to study the methyl region of the protein MSG. This 723-residue protein has been studied extensively by NMR^20,23^, but all of these studies have relied on having a deuterated samples to minimise the broadening of signals due to extensive relaxation. However, as shown in Figure 3e, by coupling the intrinsic sensitivity of methyl groups with FIDNet processing it is possible to get high quality methylTROSY like spectra for this system at a fraction of the cost and with the added bonus of simultaneously giving access to signals associated with all methyl bearing side chains. As with the HDAC8 example above, clear agreements between the expected peaks in the ILV spectrum and FID-Net processed spectrum is attained and the linewidths in these two spectra are also similar. Full overlay of the FID-Net processed spectrum of MSG and the methyl-TROSY spectrum is shown in Figure S3.

Thirdly, to push the limits of the proposed method we tested its performance on the 360 kDa α7α7 ‘half-proteasome’ from *T. acidophilum*. This protein complex has an effective rotational correlation time of approximately 120 ns at 50 °C ref^24^, though the high degree of symmetry in the complex (it is composed of 14 monomeric units that form two heptameric rings) mean that there are relatively few peaks in its spectra compared to its size, see Figure 4.

**Figure 4:**
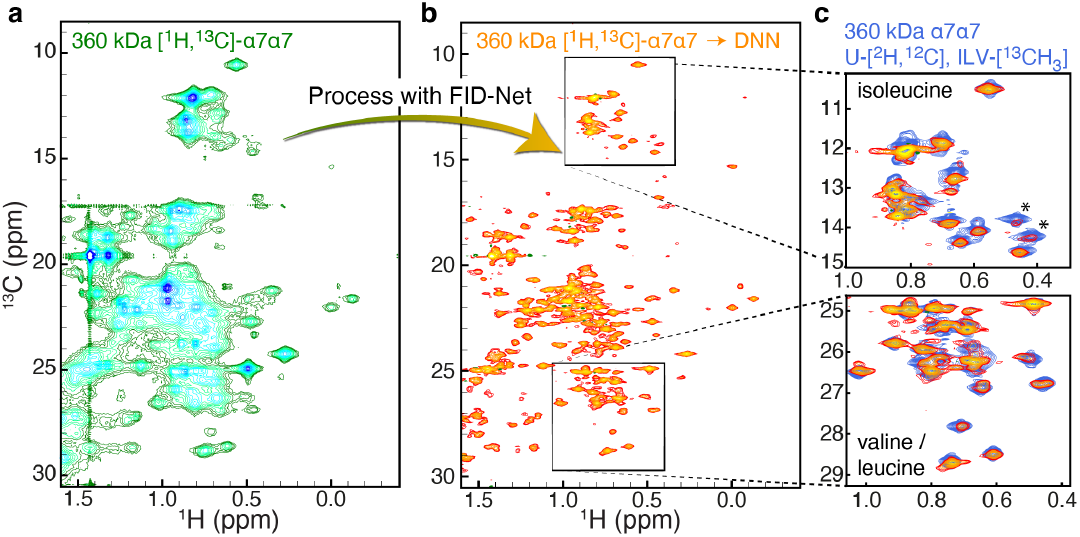
Application to a 360 kDa complex. **(a)** A ^13^C-^1^H HSQC NMR spectra of uniformly ^13^C labelled, non-deuterated, α7α7 (360 kDa) processed with a standard discrete Fourier transform. **(b)** The spectrum in a processed with the FID-Net DNNs. **(c)** Comparison of FID-Net processed HSQC spectrum in **b** (orange) with a methyl-TROSY HMQC spectrum of an ILV specifically labelled and deuterated α7α7 (blue).

In the case of the 360 kDa α7α7 complex, it is clear that a number of peaks present in the deuterated, ILV labelled sample are fairly weak in the the FID-Net decoupled spectrum (marked with an asterisk in Figure 4), this is particularly evident in the isoleucine δ1 region of the spectrum, and we therefore judge that currently systems such as the α7α7 complex is at the limit of our approach. Additional peaks in the spectrum are also clearly visible due to the presence of alanine, threonine, methionine and isoleucine ^13^C^γ2^ methyl resonances.

While for the proteins shown above the isoleucine ^13^C^δ1^, leucine ^13^C^δ1,δ2^ and valine ^13^C^γ1,γ2^ methyl group resonances can be readily compared to those obtained in ILV specifically labelled samples using traditional methyl-TROSY samples, peaks associated with alanine, threonine, methionine and isoleucine ^13^C^γ2^ methyl groups are less readily available. To verify the reliability of these additional peaks observed in uniformly ^13^C labelled samples and demonstrate the extension of the methodology to 3D spectra we recorded ^13^C^13^C-^1^H NOESY spectra of the 80 kDa MSG and as demonstrated below use this methodology to provide assignments for threonine, methionine and isoleucine side chains.

### Applications to three-dimensional NOESY spectra

Following the successful implication of deep neural networks for the production of methyl spectra of similar quality to typical deuterated methyl-TROSY spectra just from uniformly ^13^C labelled samples in a protonated background, we have applied the deep neural networks to a 3D ^13^C-HSQCNOESY-HSQC experiment acquired on a uniformly ^13^C labelled sample of MSG made in ^1^H_2_O. Despite high molecular weight of MSG, we have observed NOE cross-peaks among the inter methyl protons that were within a distance of 3.0 Å to 5.0 Å from each other, Figure 5a-5d and Figure S5. In total, approximately 312 NOE cross-peaks were observed among 170 methyl bearing residues from different regions of the protein. Simultaneously, like conventional NOESY spectra, we observe a correlation between NOE cross-peaks volume (*V*) and the distance between the proton pair (*r*), *i*.*e. V* ∝1/*r*^6^, Figure 5f. Using this experiment, we can easily connect two methyls, for example, ^13^C^δ1^ and ^13^C^γ2^ of isoleucines as shown in Figure S5a. Similarly, two geminal methyl resonances of leucine and valine can be linked using this spectrum, which is mostly present with a maximum intensity in the spectrum, Figure 5d. However, a combination of 3D ^13^C-HSQC-NOESY-HSQC with 3D HMBC-HMQC would be the most appropriate method to link the geminal methyl resonances of leucine and valine without any ambiguity^25^. Additionally, using this approach NOEs can be observed between methyl protons of all methyl bearing residues (isoleucine, leucine, valine, methionine, alanine, threonine) using one sample, which is not possible in the conventional method due to metabolic scrambling of the amino acid in selective ^13^C labelling^26^. Therefore, the deep neural network can be utilized to produce 3D ^13^C-HSQC-NOESY-HSQC experiment acquired on uniformly ^13^C labelled sample without deuteration, which results in a spectrum equivalent to 3D ^13^C-HMQC-NOESY-HMQC experiment acquired on methyl labelled sample. This approach will help the study of large proteins and complexes without deuteration.

**Figure 5:**
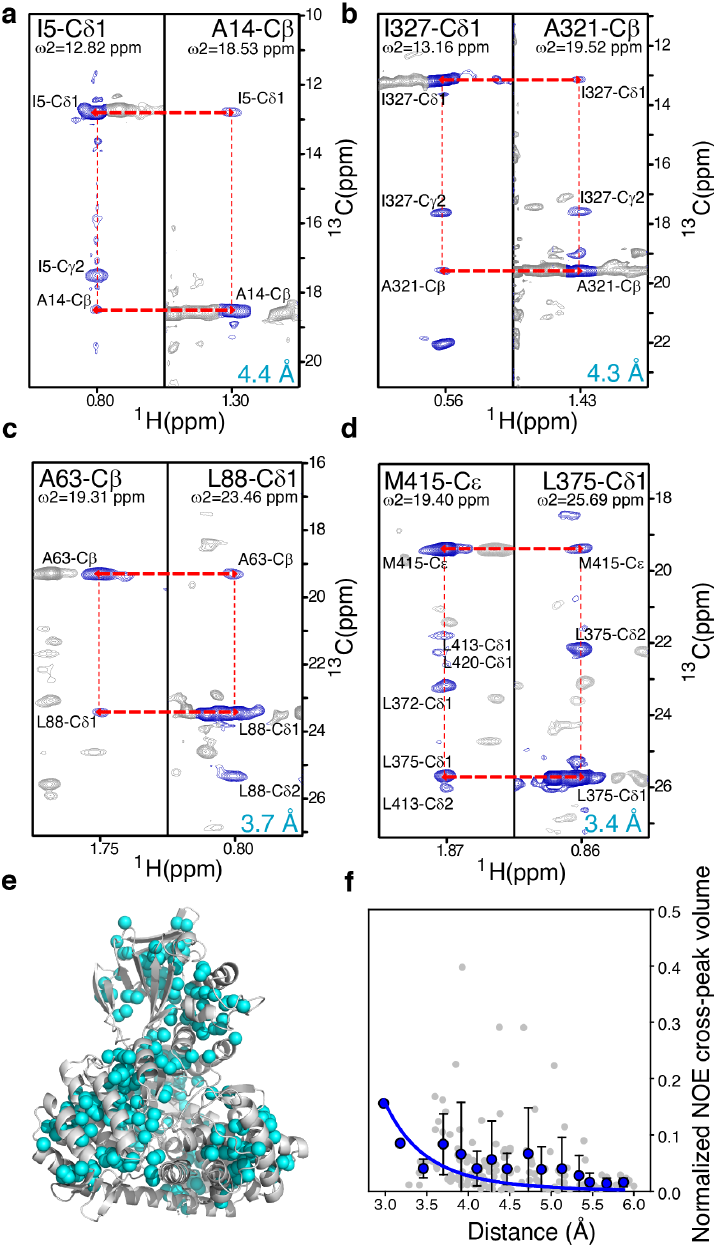
NOESY spectra of non-deuterated 80 kDa MSG. **(a)**-to-**(d)** 2D planes of the 3D ^13^C-^13^C-^1^H NOESY spectra for methyl planes of **(a)** I5-^13^C^δ1^ and A14-^13^C^β^. **(b)** I327-^13^C^δ1^ and A321-^13^C^β^. **(c)** A63-^13^C^β^ and L88-^13^C^δ1^. **(d)** M415-^13^C^ε^ and L375-^13^C^δ1^. **(e)** Methyl groups of Ile, Leu, Val, Met, Ala, and Thr showing NOE cross-peaks in 3D ^13^C-^13^C-^1^H NOESY spectra are highlighted as cyan sphere on cartoon presentation of Malate Synthase G (MSG) structure [PDB ID:1D8C]. **(f)** Normalized NOE cross-peak volumes versus interproton distances. Gray circles in the plot represent the individual data points obtained for each crosspeak. Blue circles represent the average normalized NOE cross-peaks volume over interproton-distance intervals of 0.2 Å. The blue line represents the fitted curve of NOE cross-peaks volume (V) and interproton distance (*r*) using equation *V* = C/*r*^6^, where C is a constant.

## Discussions

Being able to characterise the regulation, interactions, and dynamics of large proteins in solution is paramount to understand molecular functions. Methyl-TROSY methodologies have been one of the most important developments in biomolecular NMR over the last decades and these methods have truly paved the way for NMR becoming a complementary technique, to *e*.*g*. cryo-electron microscopy, to provide key insight on larger biomolecular complexes^27^. However, the labelling requirements for these experiments are arduous, ideally requiring perdeuteration and the use of specific precursors to introduce [^1^H,^13^C]-labelled methyl moieties in specific locations. While the resulting spectrum are of outstanding quality, the cost of such labelling is high, typically leads to lower protein yields and is inconsistent with protein production methods for many systems of interest such as eukaryotic and membrane proteins. Above, we presented an alternative method to classical methyl-TROSY NMR to characterise large proteins in solutions, which is based on uniformly protonated samples and processing with FID-Net neural networks, with which one can characterise proteins up to about 350 kDa.

The main disadvantage of the FID-Net method is that the process of peak-sharpening inevitably leads to the intrinsic peak intensity being lost. Accurately measuring peak intensities is critical in a number of NMR experiments, including diffusion and relaxation, so it is not advised to record these experiments in conjunction with FID-Net processing. However, for a large body of NMR experiments the main parameter of importance is the chemical shift as well as a reasonable estimate of the peak intensity, and in these cases we believe that FID-Net processing will prove extremely useful. Some of the uses we envisage include interaction studies, titrations, and facile chemical shift assignment of methyl peaks by either NOESY spectra (as demonstrated here for MSG) or by point mutations, which often requires several samples.

A number of approaches have previously been suggested to overcome the limitations of methyl-TROSY highlighted above, particularly when making perdeuterated samples is not possible. Recent examples include the use of delayed decoupling, as has also been used for very large complexes with molecular weights in the MDa range^28^, and optimised NMR pulse sequences to probe methionine residues in proteins with molecular weights up to 240 kDa^26^ as well as the use of local deuteration of leucine residues to probe their methyl groups in membrane and insect cell derived proteins^29^. While very powerful, these methods are limited in that they only consider a single residue type, thus restricting the number of available probes in the system. Conversely, the methodology developed here and based on processing with deep neural networks offers simultaneous access to all methyl bearing side chains in a protein, offering many more probes of biomolecular behaviour. By decoupling signals in the ^13^C dimension and sharpening them in both the ^1^H and ^13^C dimensions the resulting spectra resemble those given by perdeuterated samples with specific methyl labelling.

## Conclusion

We believe that our new methodology will significantly lower the barrier to entry for NMR of large systems. Indeed, even for well-studied systems such as MSG, while methodologies exist for obtaining methionine and threonine assignments, the processing with the FID-Net DNNs provides a straightforward method. Finally, we envisage that the idea of using DNNs for peak sharpening and simultaneous homonuclear virtual decoupling within NMR could be applied in other cases to improve spectra and that processing NMR data with DNNs, as opposed to a standard 1822 Fourier transform, will allow for many new ventures within NMR. As such we see the presented method as merely representing a “firing of the starting gun” that will pave the way for a plethora of ways to generally analysing and transforming NMR spectra with deep neural networks.

## Acknowledgements

The BBSRC (BB/R000255/1), Wellcome Trust (ref. 101569/z/13/z), and the EPSRC are acknowledged for supporting the NMR facility at University College London. Access to ultra-high field NMR spectrometers was supported by the Francis Crick Institute through provision of access to the MRC Biomedical NMR Centre and by the University of Oxford Wellcome Institutional Strategic Support Fund, the John Fell Fund, as well as the Edward Penley Abraham Cephalosporin Fund, and the Engineering and Physical Sciences Research Council (EP/R029849/1). The Francis Crick Institute receives its core funding from Cancer Research UK (FC001029), the UK Medical Research Council (FC001029), and the Wellcome Trust (FC001029). This study made use of NMRbox: National Center for Biomolecular NMR Data Processing and Analysis, a Biomedical Technology Research Resource (BTRR), which is supported by NIH grant P41GM111135 (NIGMS). Some computational aspects of this work were supported by the Francis Crick Institute (DFH) through provision of access to the Scientific Computing STP and the Crick data Analysis and Management Platform (CAMP). The Francis Crick Institute (CAMP) receives its core funding from Cancer Research UK (FC010233), the UK Medical Research Council (FC010233), and the Wellcome Trust (FC010233). For the purpose of open access, the author has applied a Creative Commons Attribution (CC BY) licence to any Author Accepted Manuscript version arising. This research is supported by the UKRI (EP/X036782/1). This paper was typeset with the bioRxiv word template by @Chrelli: www.github.com/chrelli/bioRxiv-word-template

## Author contributions

G.K. designed and trained all the DNNs, G.K. and V.K.S. produced isotope labelled samples; G.K., V.K.S., and D.F.H. performed NMR experiments; V.K.S assigned the chemical shifts of methyl spectrum. G.K. and D.F.H designed the research. All authors analysed the data, discussed the results, and wrote the paper.

## Materials and Methods

### Initial considerations about the neural networks

In the present study our aim was to develop DNNs to map ^13^C-^1^H methyl correlation spectra of large uniformly ^13^C-labelled proteins into spectra that are similar to methyl-TROSY spectra of highly deuterated proteins. Two objectives must be fulfilled to achieve this aim: (*i*) the one-bond ^13^C-^13^C homonuclear scalar couplings associated with methyl groups must be decoupled and (*ii*) the peaks must be sharpened, making them more easily resolvable, equivalent to slowing down the exponential decay of magnetisation in the time domain. It should be noted that these changes do not increase the information content in the spectrum, but they do make the information contained within the original spectra interpretable by spectroscopists.

We employ the FID-Net architecture that we have previously shown to successfully perform a number transformations on time domain NMR data, including reconstructing of non-uniformly sampled spectra^7,9,13^ and homonuclear virtual decoupling^8^. The FID-Net architecture currently only transforms a set of 1D spectra, and two separate FID-Net DNNs were therefore trained: one was optimised for spectral parameters typically encountered in the ^13^C dimension and it was trained to both decouple and sharpen signals, whereas the second FID-Net DNN was optimised for spectral parameters in the ^1^H dimension alone and was trained to only sharpen signals. A schematic illustration and summary of the effects of the neural networks is provided in Figure 1b. In both cases the networks are trained and validated exclusively on synthetic data and then tested on experimentally acquired data.

Care must be taken when determining how and to what extent signals should be sharpened using DNNs. Effectively, signals from flexible regions of proteins that already give rise to sharp could result in the presence of truncation artefacts in the spectrum. On the other hand, broad signals require significant attenuation of their relaxation to be clearly resolvable in the frequency domain. The broader the signal, the more attenuation of relaxation that is therefore desirable. To satisfy these requirements the following function is used for input 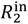 and target 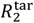 transverse relaxation rates in the training data:

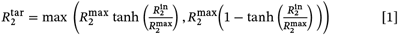

The effect of this is to make the linewidths in the target spectrum relatively uniform similar to methyl-TROSY spectra, where for the DNN relaxation rates above 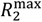 are scaled down towards it, while those below 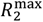 are scaled up towards it. A value of 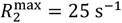 was chosen so that target spectra have linewidths similar to those observed in relatively structured residues in an ILV sample of a medium-to-large deuterated protein. Further details of the neural network architecture, training data parameters and training procedures are provided in the supplementary information.

Once trained the neural networks can easily be applied as part of processing scripts, examples of which are provided in the supplementary information. The DNNs are trained on a diverse range of NMR parameters (Table S1 and S2) and so can be used without need for further retraining and the approach can be used with standard ^1^H-^13^C HSQC or HMQC pulse sequences (vide infra).

### The network architecture and training

Two networks were trained for the study: one for removing ^13^C-^13^C couplings and sharpening spectra in the ^13^C dimension and the second purely for sharpening spectra in the ^1^H dimension (as described above). Both networks use the previously described FID-Net architecture. The input size for the ^13^C network is 1024×4 and for the ^1^H network is 512×4. As is the case with previous FID-Net architectures, the networks consist of a series of stacked residual units, wherein each residual unit consists of dilated convolutional layers with kernel size 8×4. The filters are activated by either sigmoidal (50%) or tangent (50%) functions. The results of the activations are then multiplied and passed through another convolutional layer with kernel size 8×4. As described previously, the output from each layer is combined to give the final output and also added to the input for the residual unit to form the input for the next layer. For the ^13^C network, the dilations employed are cycled through the values: 1,2,4,6,8,10,12,14,16,20,24,28,32,40,48,56,64, and there are 128 filters for each convolutional layer. For the ^1^H network the dilations employed are 1,2,4,6,8,10,12,14,16,20,24,28,32, and there are 64 filters per convolutional layer.

For each network 500,000 test planes were created for training and 50,000 for cross-validation using the parameters given in Tables S1 and S2. The models were developed and trained using the Tensorflow library^30^ with the Keras-front end^31^. The cost function used to train the networks is the mean squared error in the frequency domain between the spectrum produced by the DNN and the target spectrum wherein the linewidth of peaks are set according to the R_2_ scaling described above and for the ^13^C network the scalar coupling is removed. The RMSprop optimizer^32^ is used in training. For both networks the learning rate was initially set to 10^-4^ until the validation loss value plateaus and is then reduced to 10^-5^ until it plateaus again where training is then ended.

### Cross-validations using synthetic data

Once trained, to cross validate the performance of the two networks we use synthetically generated spectra. Rather than using arbitrary spectra as was done for training the networks, we attempt to generate realistic ^13^C-^1^H correlation maps for uniformly labelled ^13^C proteins using chemical shift statistics from the BMRB^18^. In addition to containing terminal ^13^C moieties that give rise to doublets in the spectra as a result of a single ^13^C-^13^C scalar coupling these spectra also contain multiplets due to moieties that have multiple ^13^C-^13^C scalar couplings (though these are usually at higher ^1^H frequencies as is observed in real spectra).

We generate two hundred synthetic spectra in total: the first hundred spectra are chosen to have features similar to a ^13^C-^1^H spectra of a protein with a similar size to HDAC whilst the second hundred are chosen to have features for a protein with a similar size to MSG. Example spectra are shown in Figure 2A and 2B. The following parameters are used to generate the synthetic spectra for cross-validation:

| Parameter | HDAC-like spectra | MSG-like spectra |
| --- | --- | --- |
| Number of signals | 275 | 600 |
| Larmor Frequency (MHz) | $\in (600,700,800,950)$ | $\in (600,700,800,950)$ |
| $^1\text{H}$ SW (Hz) | 2000 - 5000 | 2000 - 5000 |
| $^{13}\text{C}$ SW (Hz) | 2000 - 5000 | 2000 - 5000 |
| $^1J_{\text{CC}}$ (Hz) | 34 (2) | 34 (2) |
| $R_2^{(1)}$ ($\text{s}^{-1}$) ($^1\text{H}$ dim) | 50 (10) | 60 (15) |
| $R_2^{(2)}$ ($\text{s}^{-1}$) ($^{13}\text{C}$ dim) | 50 (10) | 60 (10) |

Where values are given as a range, for each spectrum the true value is taken as a random value from a uniform distribution of the range. Where values are given as a number followed by a bracketed number, for each signal the value used is randomly chosen from a normal distribution centred on the first number and with a standard deviation given by the bracketed value. For all synthetic spectra, half of the peaks are chosen as resulting from a methyl moiety, while the other half come from non-methyl ^13^C-^1^H moieties, *i*.*e*., leading to triplets in the ^13^C dimension.

Once the synthetic spectra are made they are processed using the deep neural network pipeline. Visual inspection of the transformed spectra suggests that the method is effective at decoupling and sharpening spectra, such that they can be interpreted more easily. We also note that where multiple ^13^C-^13^C couplings are present the network will remove just one of the couplings such that triplets become doublets. To evaluate the performance of the pipeline quantitatively, the transformed spectra are peak picked using the built-in peak picker in NMRPIPE. The results are compared to the known ground-truth values for peak positions (assuming no couplings were present in the spectra).

We focus on two key parameters in this evaluation: the number of true positive picked peaks and the number of false positive picked peaks. Given the difficulty in picking peaks from crowded regions automatically, and our aim here to evaluate the performance of our alternative pipeline for analysing spectra we focus attention on isolated peaks where the performance of the peak picker is robust. Here, we define an isolated peak as any ground truth peak where the minimum distance from any other ground truth peak is greater than or equal to 0.06 ^1^H ppm (0.24 ^13^C ppm). Furthermore, we ignore peaks that are part of doublets and that are more than 0.25 ^1^H ppm from a target peak originating from a methyl moiety. Once peaks in the FID-Net processed spectrum have been picked they are matched with the closest peak in the target peak list. Each peak can only be matched with a single target peak and we match in descending order of distance between picked peaks and target peaks until there are no picked peaks within the minimum distance to a target peak (set at 0.05 ^1^H ppm) or there are no picked or target peaks left. This process is then repeated for all synthetic spectra.

The true positive rate is defined as the percentage of peaks where we see a clear correspondence between a picked and target peak. The false positive rate is defined as the rate at which a peak identified in the FID-Net processed spectrum does not correspond with any peaks in the actual spectrum. Given the limitations of peak pickers, the true and false positive rates here likely represent a lower bound on the performance of the FID-Net processing pipeline.

### Sample preparations

#### General isotopic labelling

In this study, two different types of labelled protein samples were used for acquisition of the NMR data. (1) uniformly ^13^C,^15^N labelled sample made in ^1^H_2_O. (2) methyl Ile-^13^C^δ1^,^1^H^δ1^, Leu-^13^C^δ1^,^1^H^δ1^/^13^C^δ2^,^1^H^δ2^, and Val-^13^C^γ1^,^1^H^γ1^/^13^C^γ2^,^1^H^γ2^ labelled sample made in deuterated background. To express uniformly ^13^C,^15^N labelled proteins, we used M9 media that was made with ^1^H_2_O and supplemented with 1 g/L [^1^H,^15^N]-ammonium chloride and 3 g/L of [^1^H,^13^C]-glucose as the sole nitrogen and carbon sources. While for expression of methyl Ile-^13^C^δ1^,^1^H^δ1^, Leu-^13^C^δ1^,^1^H^δ1^/^13^C^δ2^,^1^H^δ2^, and Val-^13^C^γ1^,^1^H^γ1^/^13^C^γ2^,^1^H^γ2^ labelled proteins, we used ^2^H_2_O M9 media supplemented with 1 g/L [^1^H,^15^N]-ammonium chloride and 3 g/L of [^2^H,^12^C]-glucose as the sole nitrogen and carbon sources. Methyl labelling was achieved by the addition of 60 mg/L alpha-ketobutyric acid [U-^12^C/^2^H, methyl-^13^CH_3_] for labelling of isoleucines, and 90 mg/L αketoisovaleric acid [U-^12^C/^2^H, methyl-(^13^CH_3_,^12^CD_3_)] for labelling of valine and of leucine methyl groups. These precursors were added one hour prior to induction.

#### Expression and purification of Histone de-acetylase 8 (HDAC8)

The Human HDAC8 construct described by Vannini et al. with a C-terminal 6X-histidine tag in ampicillin-resistant pET21b expression vector was transformed in BL21(λDE3) *E. coli* cells for protein expression^19,33^. A single colony from the transformed plate was inoculated in 10 ml of LB media supplemented with ampicillin (100 µg/ml) at 37°C. Once the LB culture reached an OD600 between 0.8 and 1.0, it was used to inoculate a 50 ml M9 minimal media pre-culture. This M9 pre-culture was used to inoculate 1 L of M9 media and grown at 37°C to OD600 ≈ 0.8. HDAC8 expression was induced for >16 hour with 0.5mM IPTG and 200 µM of ZnCl_2_ at 21°C. The cell pellet, collected by centrifugation, was re-suspended in lysis buffer containing 50 mM Tris-HCl pH 8.0, 3mM MgCl2, 500 mM KCl, 10 mM imidazole, 5% glycerol, and 10 mM β-mercaptoethanol. Later, sonication was performed to lyse the cells after addition of small amounts of DNAse, lysozyme, protease inhibitors tablets (1 tablet per 50 ml, Roche), and 0.25 % IGEPAL. The supernatant fraction of lysate after centrifugation at 18000 rpm for one hour was purified by Ni-NTA affinity chromatography using a linear imidazole gradient (10‒250 mM) in lysis buffer. Further, a size-exclusion chromatography on Superdex-75 column (GE Healthcare) was carried out in buffer containing 50 mM Tris-HCl pH 8.0, 150 mM KCl, 1 mM TCEP, and 5% glycerol. Fractions containing purified HDAC8 was pooled together and concentrated by 10 kDa cut off Amicon (Millipore) ultra-filtration membranes. The concentrated sample was buffer exchanged into NMR-buffer (50 mM K_2_HPO_4_ pH 8.0, 30 mM KCl, 4 mM DTT, and 1 mM NaN_3_) for NMR data acquisition.

#### Expression and purification of Malate synthase G (MSG)

A small adjustment was made to the methods previously described for producing isotopically labelled MSG^23,25,34^. Briefly, to produce the MSG protein, BL21 (DE3) *E. coli* cells were transformed with a kanamycin-resistant pET28a vector containing MSG gene with a C-terminal 6Xhistidine tag. The protein expression protocol for MSG is same as HDAC8 up to induction. MSG expression was induced for >16 hour with 1mM IPTG at 21°C. The cell pellet was collected by centrifugation and re-suspended in lysis buffer containing 20 mM Tris-HCl pH 7.8, 300 mM NaCl, 10 mM imidazole, and 10 mM β-mercaptoethanol. Like HDAC8, the protein purification protocol for MSG is same up to Ni-NTA affinity chromatography. The fractions containing MSG from Ni-NTA affinity chromatography were further purified by size exclusion chromatography using a Superdex-200 column (GE Healthcare) in buffer containing 20 mM Sodium phosphate pH 7.1, 5mM dithiothreitol. After gel filtration, the fractions containing pure protein were pooled, concentrated, and buffer exchanged into NMR-buffer (20 mM Sodium phosphate buffer pH 7.1, 5 mM DTT, 20 mM MgCl_2_, 1 mM NaN_3_) for NMR data acquisition using 30 kDa cut off Amicon (Millipore) ultra-filtration membranes.

#### Expression and purification of α-subunit complex (α7α7) of proteasome from T. acidophilum

In order to express the α-subunit complex (α7α7) proteasome from *T. acidophilum*, the αWT clone with N-terminal Histidine tag and a TEV protease site was transformed into BL21 (λDE3) E. coli cells^25,35^. The protein expression protocol for this α-subunit complex is same as MSG up to induction. The α-subunit complex culture was induced at OD600 ≈ 0.9 with 1mM IPTG at 37°C for 5 hours. Afterward, we lysed the cells with sonication in a lysis buffer (50 mM NaH2PO4 pH 8.0, 0.2 M NaCl, 10 mM imidazole) and purified them using Ni-NTA chromatography as described above in purification sections of HDAC8 and MSG. After Ni-NTA affinity chromatography, TEV protease was introduced to cleave the 6X-histidine tag before dialyzing the protein against 2 L of dialysis buffer (50 mM Tris-HCl pH 8.0, 1 mM EDTA, 5 mM β-mercaptoethanol) overnight at 4°C. The TEV cleavage of the protein was followed by another Ni-NTA affinity chromatography to eliminate the histidine tag and un-cleaved protein. Afterward, a size exclusion chromatography was performed using a Superdex 200 column (GE Healthcare) in a buffer containing 50 mM NaH_2_PO_4_ pH 7.5, and 100 mM NaCl. The fractions containing pure protein was concentrated and buffer exchanged in NMR buffer (20 mM potassium phosphate pH 6.8, 50 mM NaCl, 1 mM EDTA, 2 mM DTT, 0.03% NaN_3_) for NMR data acquisition using 30 kDa cut off Amicon (Millipore) ultra-filtration membranes.

### NMR acquisitions

#### Three-dimensional NOESY spectra

The 3D HSQC-NOESY-HSQC NMR experiment on MSG was performed on a ∼400 µM MSG sample on a Bruker 950 MHz Avance HD spectrometer equipped with Z-gradient triple-resonance TCI cryoprobe. The data was acquired with 1024, 142, and 124 complex points in ^1^H, ^13^C_HSQC_, and ^13^C_NOESY_ dimensions, respectively, with spectral widths of 15243.9 Hz (^1^H), 6666.7 Hz (^13^C), and 6666.7 Hz (^13^C). Eight scans were collected per increment with a recycle delay of 1 s and the mixing time was 60 ms.

#### Two-dimensional ^13^C-^1^H correlation spectra

The 2D HSQC spectrum of HDAC8 used as input for FID-Net was recorded on a uniformly [^13^C,^15^N]-labelled sample using a standard pulse programme with presaturation on a Bruker 700 MHz Avance III spectrometer equipped with Z-gradient triple-resonance TCI cryoprobe. The data was acquired with 1024 and 256 complex points in the ^1^H and ^13^C dimensions, respectively, with spectral widths of 10000 Hz and 5000 Hz. 16 scans were obtained per individual FID. 2D Methyl-TROSY spectrum of HDAC8 used for cross-validation was recorded on an ILVM specifically labelled sample using a standard pulse programme on a Bruker 800 MHz Avance III spectrometer equipped with Z-gradient triple-resonance TCI cryoprobe. The data was acquired with 1024 and 256 complex points in the ^1^H and ^13^C dimensions, respectively, with spectral widths of 12500 Hz and 4500 Hz. 4 scans were obtained per individual FID.

The 2D HSQC spectrum of MSG used as input for FID-Net was recorded on a uniformly [^13^C,^15^N]-labelled sample using a standard pulse programme with presaturation on a Bruker 800 MHz Avance III spectrometer equipped with Z-gradient triple-resonance TCI cryoprobe. The data was acquired with 1024 and 256 complex points in the ^1^H and ^13^C dimensions, respectively, with spectral widths of 12500 Hz and 5000 Hz. 16 scans were obtained per individual FID. 2D Methyl-TROSY spectrum of MSG used for cross-validation was recorded on an ILV specifically labelled sample using a standard pulse programme on a Bruker 800 MHz Avance III spectrometer equipped with Z-gradient triple-resonance TCI cryoprobe. The data was acquired with 1024 and 192 complex points in the ^1^H and ^13^C dimensions, respectively, with spectral widths of 10500 Hz and 5000 Hz. 16 scans were obtained per individual FID.

The 2D HSQC spectrum of α7α7 used as input for FID-Net was recorded on a uniformly [^13^C,^15^N]-labelled sample using a standard pulse programme with presaturation on a Bruker 950 MHz Avance HD spectrometer equipped with Z-gradient triple-resonance TCI cryoprobe. The data was acquired with 1024 and 256 complex points in the ^1^H and ^13^C dimensions, respectively, with spectral widths of 15200 Hz and 5263 Hz. 80 scans were obtained per individual FID. 2D Methyl-TROSY spectrum of MSG used for cross-validation was recorded on an ILV specifically labelled sample using a standard pulse programme on a Bruker 800 MHz Avance III spectrometer equipped with Z-gradient triple-resonance TCI cryoprobe. The data was acquired with 768 and 132 complex points in the ^1^H and ^13^C dimensions, respectively, with spectral widths of 12000 Hz and 4100 Hz. 16 scans were obtained per individual FID.

## Data processing

All experimental NMR spectra were processed with NMRPIPE^36^ or using the python libraries NMRGLUE^37^ and NUMPY.

## Code Availability

Code (python) for using the networks described here (including pretrained networks and examples) is available on GitHub: https://github.com/gogulan-k/FID-Net

**Table S1:**
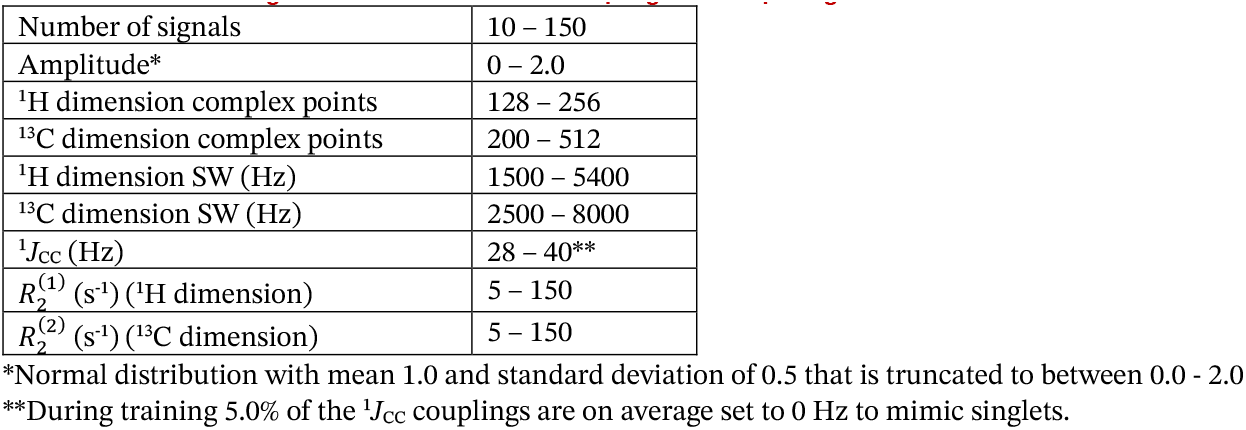
Parameter ranges used to train the ^13^C decoupling and sharpening network.

|  |  |
| --- | --- |
| Number of signals | 10 – 150 |
| Amplitude* | 0 – 2.0 |
| $^1\text{H}$ dimension complex points | 128 – 256 |
| $^{13}\text{C}$ dimension complex points | 200 – 512 |
| $^1\text{H}$ dimension SW (Hz) | 1500 – 5400 |
| $^{13}\text{C}$ dimension SW (Hz) | 2500 – 8000 |
| $^1J_{\text{CC}}$ (Hz) | 28 – 40** |
| $R_2^{(1)}$ ($\text{s}^{-1}$) ($^1\text{H}$ dimension) | 5 – 150 |
| $R_2^{(2)}$ ($\text{s}^{-1}$) ($^{13}\text{C}$ dimension) | 5 – 150 |

\*Normal distribution with mean 1.0 and standard deviation of 0.5 that is truncated to between 0.0 - 2.0
\*\*During training 5.0% of the $^1J_{\text{CC}}$ couplings are on average set to 0 Hz to mimic singlets.

**Table S2:**
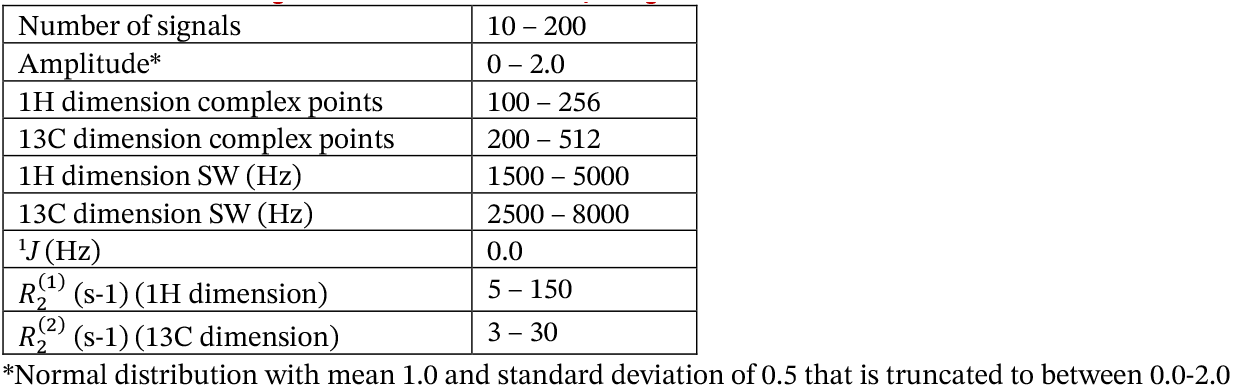
Parameter ranges used to train the ^1^H sharpening network.

**Figure S1:**
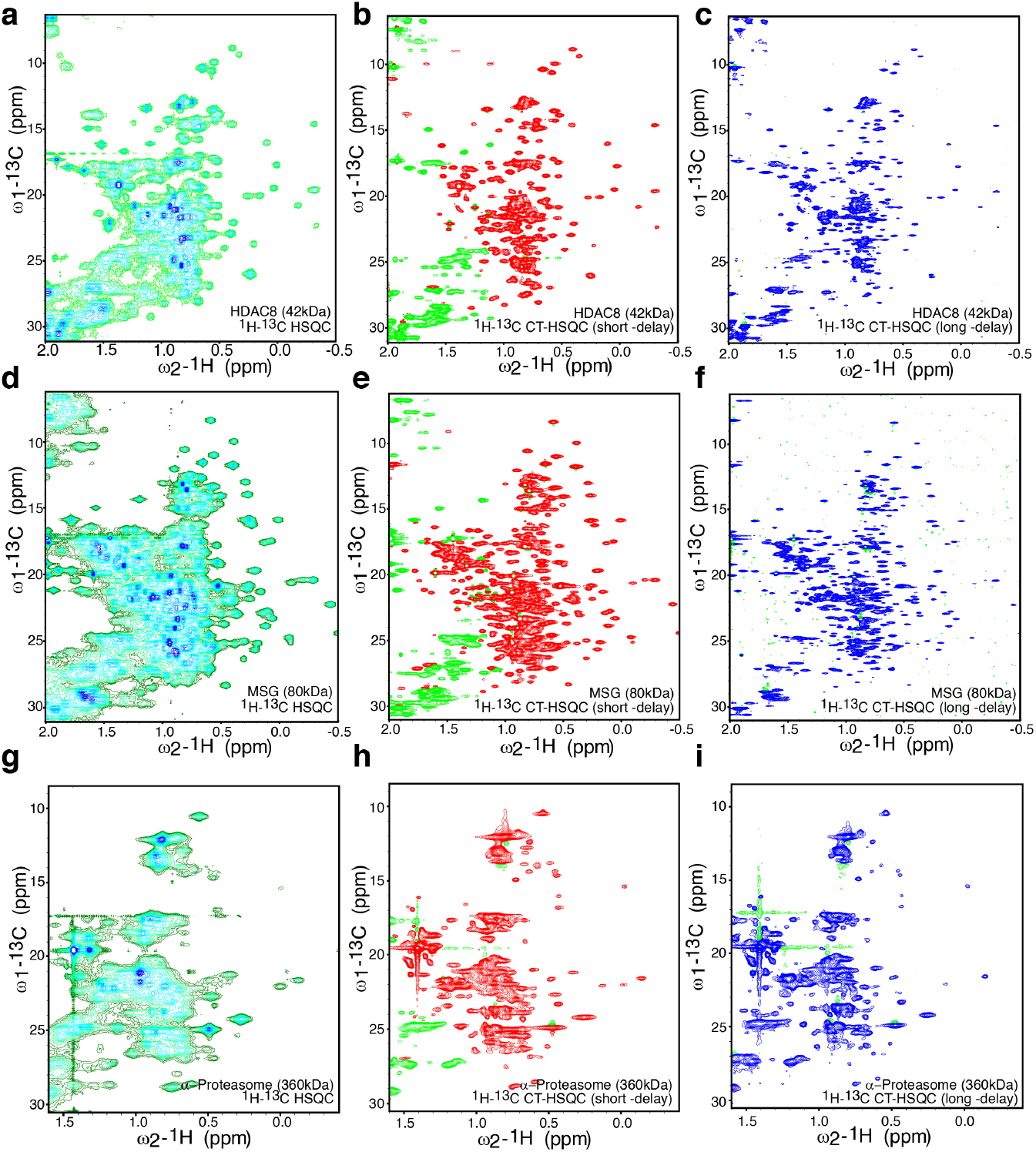
Comparison of ^13^C HSQC and 13C-Constant Time (CT)-HSQC spectra of uniformly ^13^C labelled, non-deuterated HDAC8, MSG, and α-proteasome. **(a), (d)**, and **(g)** are the ^13^C-HSQC spectra of HDAC8, MSG, and α7α7-proteasome, respectively. **(b), (e)**, and **(h)** are the ^13^C-constant-time (CT) HSQC spectra, acquired with constant-time period of 27 ms, of HDAC8, MSG, and α-proteasome, respectively. **(c), (f)**, and **(i)** are the ^13^C-CT-HSQC spectra, acquired with constant-time of 54 ms, of HDAC8, MSG, and α7α7-proteasome, respectively.

**Figure S2:**
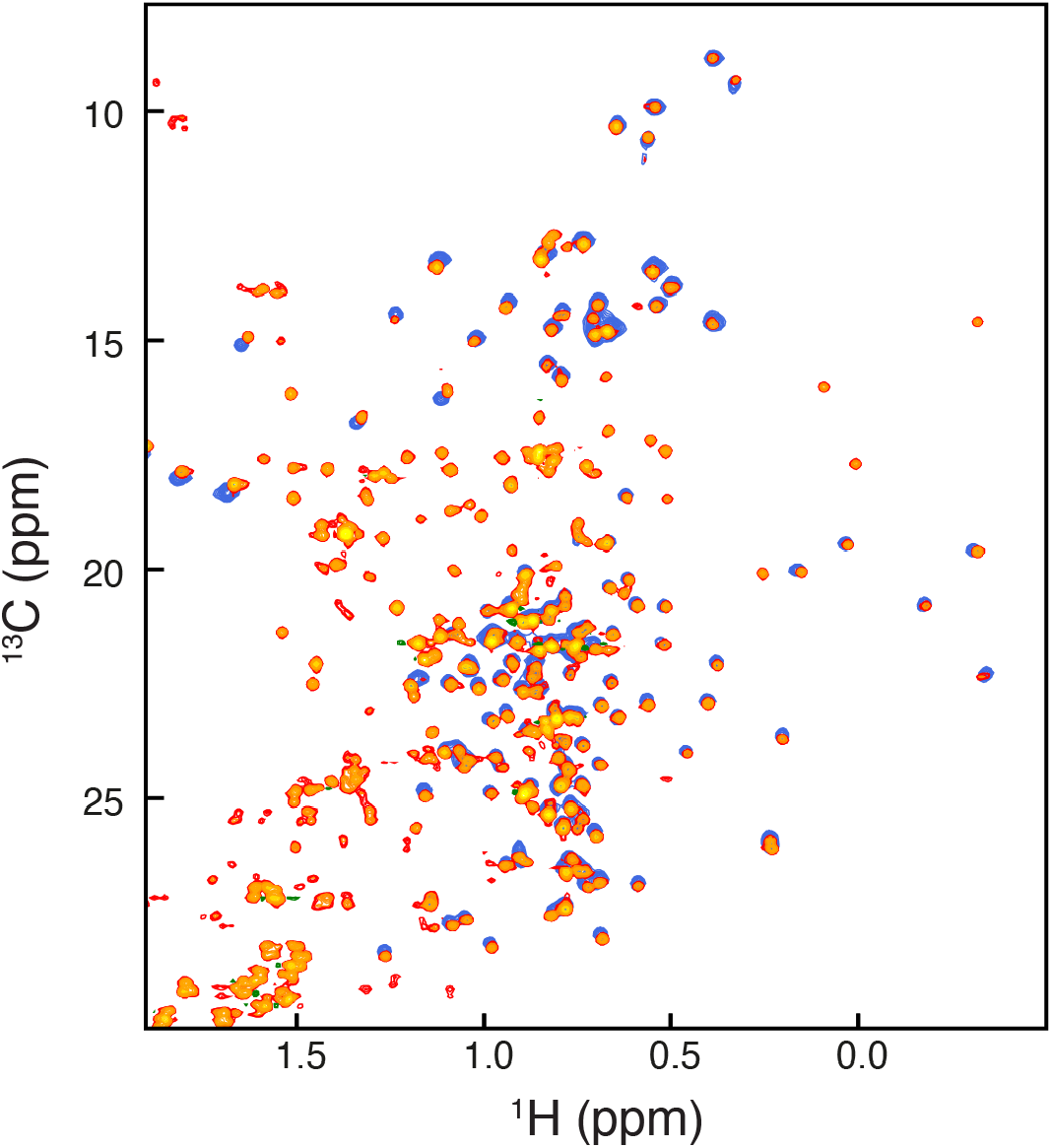
Comparison of FID-Net processed HSQC spectrum in (orange) of the 42 kDa HDAC8 (**Figure 3b**) with a methyl-TROSY HMQC spectrum of an ILVM specifically labelled and deuterated HDAC8 (blue). Small shifts are due to two-bond (^13^C) and three-bond (^1^H) deuterium isotope shifts that are different for Isoleucine (2 gamma protons), Leucine (1 gamma proton), Valine (1 beta proton), and Methionine (0 delta protons).

**Figure S3:**
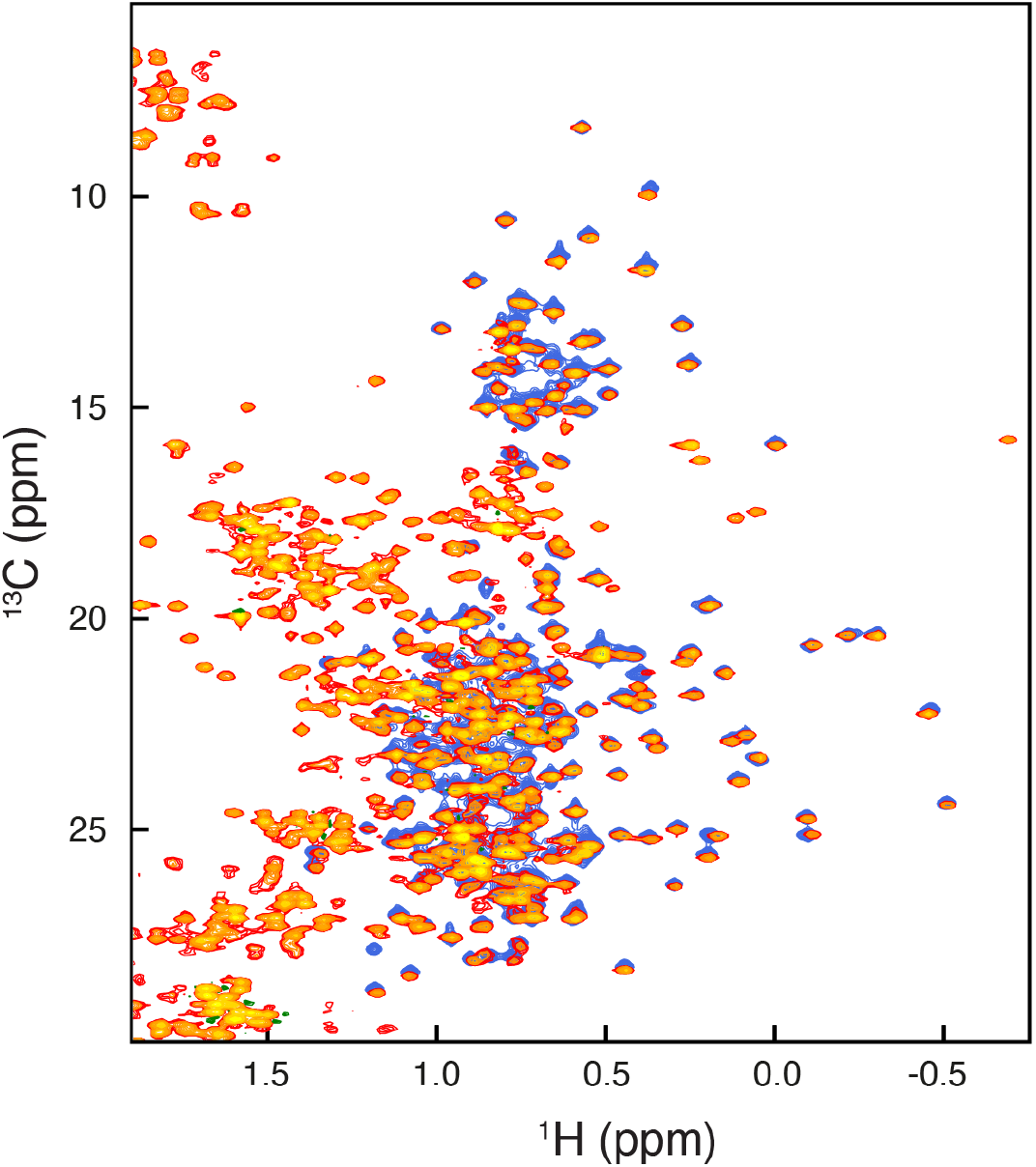
Comparison of FID-Net processed HSQC spectrum in (orange) of the 80 kDa MSG (**Figure 3e**) with a methyl-TROSY HMQC spectrum of an ILV specifically labelled and deuterated MSG (blue). Small shifts are due to two-bond (^13^C) and three-bond (^1^H) deuterium isotope shifts that are different for Isoleucine (2 gamma protons) and Leucine (1 gamma proton) and Valine (1 beta proton).

**Figure S4:**
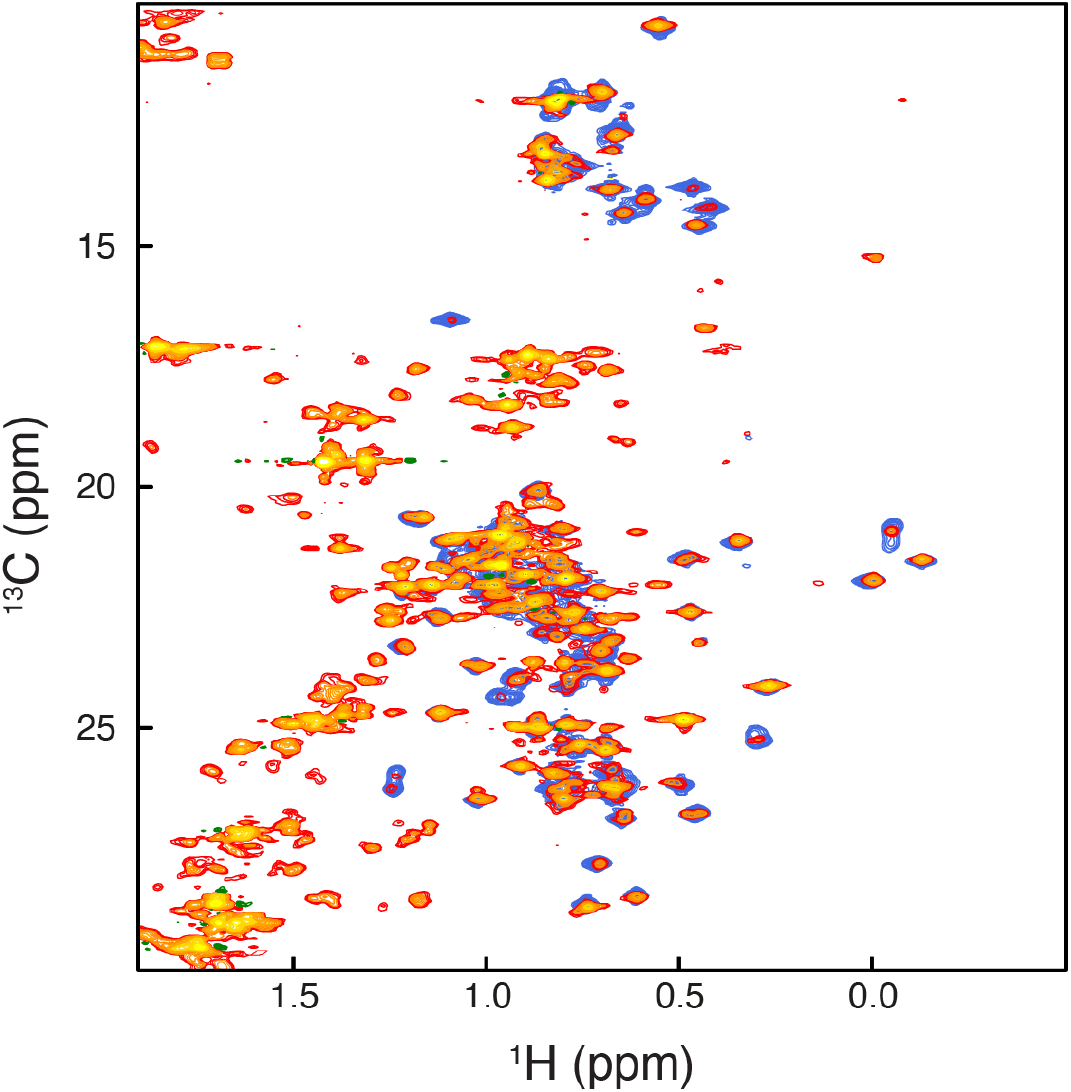
Comparison of FID-Net processed HSQC spectrum in (orange) of the 360 kDa α7α7 (**Figure 4b**) with a methyl-TROSY HMQC spectrum of an ILV specifically labelled and deuterated α7α7 (blue). Small shifts are due to two-bond (^13^C) and three-bond (^1^H) deuterium isotope shifts that are different for Isoleucine (2 gamma protons) and Leucine (1 gamma proton) and Valine (1 beta proton).

**Figure S5:**
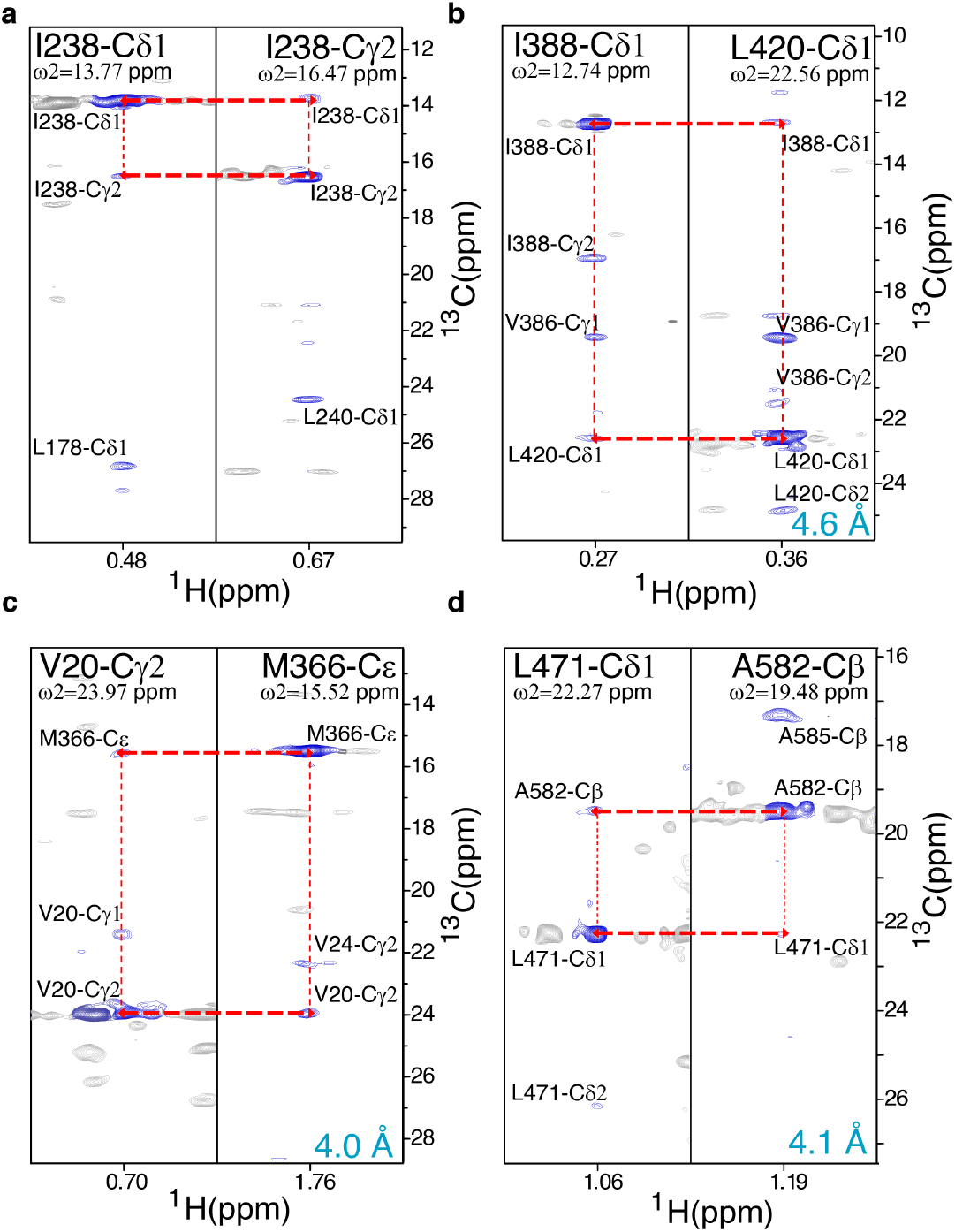
NOESY spectra of uniformly ^13^C labelled, non-deuterated 80 kDa MSG. **(a)**-to-**(d)** 2D planes of the 3D ^13^C-^13^C-^1^H NOESY spectra for methyl planes of **(a)** I238-^13^C^δ1^ and I238-^13^Cγ^**2**^. **(b)** I388-^13^C^δ1^ and L420-^13^C^δ1^. **(c)** V20-^13^Cγ^**2**^ and M366-^13^C^ε^. **(d)** L471-^13^C^δ1^ and A582-^13^C^**β**^.

## References

1. Tugarinov, V., Hwang, P. M., Ollerenshaw, J. E. & Kay, L. E. Cross-Correlated Relaxation Enhanced ^1H−13^C NMR Spectroscopy of Methyl Groups in Very High Molecular Weight Proteins and Protein Complexes. J. Am. Chem. Soc. 125, 10420–10428 (2003).

2. Tugarinov, V. & Kay, L. E. An isotope labeling strategy for methyl TROSY spectroscopy. J Biomol NMR 28, 165–172 (2004).

3. Boisbouvier, J. & Kay, L. E. Advanced isotopic labeling for the NMR investigation of challenging proteins and nucleic acids. J. Biomol. NMR 71, 115–117 (2018).

4. Jumper, J. et al. Highly accurate protein structure prediction with AlphaFold. Nature 596, 583–589 (2021).

5. Baek, M. et al. Accurate prediction of protein structures and interactions using a three-track neural network. Science. 373, 871–876 (2021).

6. Birhane, A., Kasirzadeh, A., Leslie, D. & Wachter, S. Science in the age of large language models. Nat. Rev. Phys. 5, 277–280 (2023).

7. Karunanithy, G. & Hansen, D. F. FID-Net: A versatile deep neural network architecture for NMR spectral reconstruction and virtual decoupling. J. Biomol. NMR 75, 179–191 (2021).

8. Karunanithy, G., Mackenzie, H. W. & Hansen, D. F. Virtual Homonuclear Decoupling in Direct Detection Nuclear Magnetic Resonance Experiments Using Deep Neural Networks. J. Am. Chem. Soc. 143, 16935–16942 (2021).

9. Qu, X. et al. Accelerated Nuclear Magnetic Resonance Spectroscopy with Deep Learning. Angew. Chemie 132, 10383–10386 (2020).

10. Karunanithy, G., Yuwen, T., Kay, L. E. & Hansen, D. F. Towards autonomous analysis of chemical exchange saturation transfer experiments using deep neural networks. J. Biomol. NMR 76, 75–86 (2022).

11. Li, D.-W., Hansen, A. L., Yuan, C., Bruschweiler-Li, L. & Brüschweiler, R. DEEP picker is a deep neural network for accurate deconvolution of complex two-dimensional NMR spectra. Nat. Commun. 12, 5229 (2021).

12. Worswick, S. G., Spencer, J. A., Jeschke, G. & Kuprov, I. Deep neural network processing of DEER data. Sci. Adv. 4, eaat5218 (2018).

13. Hansen, D. F. Using Deep Neural Networks to Reconstruct Non-uniformly Sampled NMR Spectra. J. Biomol. NMR 73, 577–585 (2019).

14. Klukowski, P. et al. NMRNet: a deep learning approach to automated peak picking of protein NMR spectra. Bioinformatics 34, 2590–2597 (2018).

15. Grattan-Guinness, I. Joseph Fourier, Théorie analytique de la chaleur (1822). in Landmark Writings in Western Mathematics 1640-1940 354–365 (Elsevier, 2005). doi:10.1016/B978-044450871-3/50107-8.

16. Santoro, J. & King, G. C. A constant-time 2D overbodenhausen experiment for inverse correlation of isotopically enriched species. J. Magn. Reson. 97, 202–207 (1992).

17. Vuister, G. W. & Bax, A. Resolution enhancement and spectral editing of uniformly 13C-enriched proteins by homonuclear broadband ^13^C decoupling. J. Magn. Reson. 98, 428–435 (1992).

18. Ulrich, E. L. et al. BioMagResBank. Nucleic Acids Res. 36, D402–D408 (2008).

19. Werbeck, N. D. et al. A distal regulatory region of a class I human histone deacetylase. Nat. Commun. 11, 3841 (2020).

20. Tugarinov, V. & Kay, L. E. Ile, Leu, and Val methyl assignments of the 723-residue malate synthase G using a new labeling strategy and novel NMR methods. J. Am. Chem. Soc. 125, 13868–78 (2003).

21. Kerfah, R., Plevin, M. J., Sounier, R., Gans, P. & Boisbouvier, J. Methyl-specific isotopic labeling: a molecular tool box for solution NMR studies of large proteins. Curr. Opin. Struct. Biol. 32, 113–122 (2015).

22. Hansen, P. Isotope effects on chemical shifts of proteins and peptides. Mag. Res. Chem. 38, 1–10 (2000).

23. Tugarinov, V., Muhandiram, R., Ayed, A. & Kay, L. E. Four-Dimensional NMR Spectroscopy of a 723-Residue Protein: Chemical Shift Assignments and Secondary Structure of Malate Synthase G. J. Am. Chem. Soc. 124, 10025–10035 (2002).

24. Sprangers, R. & Kay, L. E. Quantitative dynamics and binding studies of the 20S proteasome by NMR. Nature 445, 618–622 (2007).

25. Siemons, L., Mackenzie, H. W., Shukla, V. K. & Hansen, D. F. Intra-residue methyl–methyl correlations for valine and leucine residues in large proteins from a 3D-HMBC-HMQC experiment. J. Biomol. NMR 73, 749–757 (2019).

26. Rößler, P., Mathieu, D. & Gossert, A. D. Enabling NMR Studies of High Molecular Weight Systems Without the Need for Deuteration: The XL-ALSOFAST Experiment with Delayed Decoupling. Angew. Chemie Int. Ed. 59, 19329–19337 (2020).

27. Vahidi, S. et al. An allosteric switch regulates Mycobacterium tuberculosis ClpP1P2 protease function as established by cryo-EM and methyl-TROSY NMR. Proc. Natl. Acad. Sci. 117, 5895–5906 (2020).

28. Bolik-Coulon, N. et al. Less is more: A simple methyl-TROSY based pulse scheme offers improved sensitivity in applications to high molecular weight complexes. J. Magn. Reson. 346, 107326 (2023).

29. Dubey, A. et al. Local Deuteration Enables NMR Observation of Methyl Groups in Proteins from Eukaryotic and Cell-Free Expression Systems. Angew. Chemie Int. Ed. 60, 13783–13787 (2021).

30. Abadi, M. et al. TensorFlow: Large-scale machine learning on heterogeneous systems. (2015).

31. Chollet, F. and others. Keras. (2015). Available at https://github.com/fchollet/keras

32. Tieleman, T. & Hinton, G. Lecture 6.5-rmsprop: Divide the Gradient by a Running Average of Its Recent Magnitude. COURSERA: Neural Networks for Machine Learning. (2012).

33. Vannini, A. et al. Substrate binding to histone deacetylases as shown by the crystal structure of the HDAC8-substrate complex. EMBO Rep 8, 879–884 (2007).

34. Pritchard, R. B. & Hansen, D. F. Characterising side chains in large proteins by protonless ^13^C-detected NMR spectroscopy. Nat. Commun. 10, 1747 (2019).

35. Tugarinov, V., Sprangers, R. & Kay, L. E. Probing Side-Chain Dynamics in the Proteasome by Relaxation Violated Coherence Transfer NMR Spectroscopy. J. Am. Chem. Soc. 129, 1743–1750 (2007).

36. Delaglio, F. et al. Nmrpipe - a Multidimensional Spectral Processing System Based on Unix Pipes. J. Biomol. Nmr 6, 277–293 (1995).

37. Helmus, J. J. & Jaroniec, C. P. Nmrglue: an open source Python package for the analysis of multidimensional NMR data. J. Biomol. NMR 55, 355–367 (2013).

